# Computational Design and Biophysical Validation of Macrocyclic Peptides as Inhibitors of SLIT2/ROBO1 interaction

**DOI:** 10.1101/2025.10.26.684696

**Authors:** Somaya A. Abdel-Rahman, Maxence Delaunay, Tâp Ha-Duong, Moustafa Gabr

## Abstract

The SLIT2/ROBO1 signaling axis regulates cellular migration and angiogenesis but also contributes to tumor progression and immune evasion in glioblastoma. Targeting this pathway with small molecules or antibodies remains challenging due to the shallow and extended nature of the SLIT2/ROBO1 interface. Here, we report the first computational design and experimental validation of macrocyclic peptides that inhibit SLIT2/ROBO1 binding. Twenty peptides were generated through a structure-guided interface mapping approach (Des3PI 2.0) and ranked using a contact-based scoring function. The top candidates were synthesized and evaluated using time-resolved fluorescence resonance energy transfer (TR-FRET) and biolayer interferometry (BLI) assays. Among the SLIT2-targeting peptides, **SP4** and **SP3** showed the most pronounced inhibition in TR-FRET and BLI, confirming direct binding to the SLIT2/ROBO1 interface. The lead peptide **SP4** also demonstrated favorable in vitro pharmacokinetic properties, including strong stability in simulated intestinal fluid, high plasma integrity, and moderate metabolic stability in rat liver microsomes. Collectively, this work establishes a computational-to-experimental pipeline for discovering macrocyclic peptides that disrupt challenging protein-protein interactions and provides a foundation for developing next-generation SLIT2/ROBO1 modulators for cancer and neuroimmune disorders.

**Graphical Abstract:** 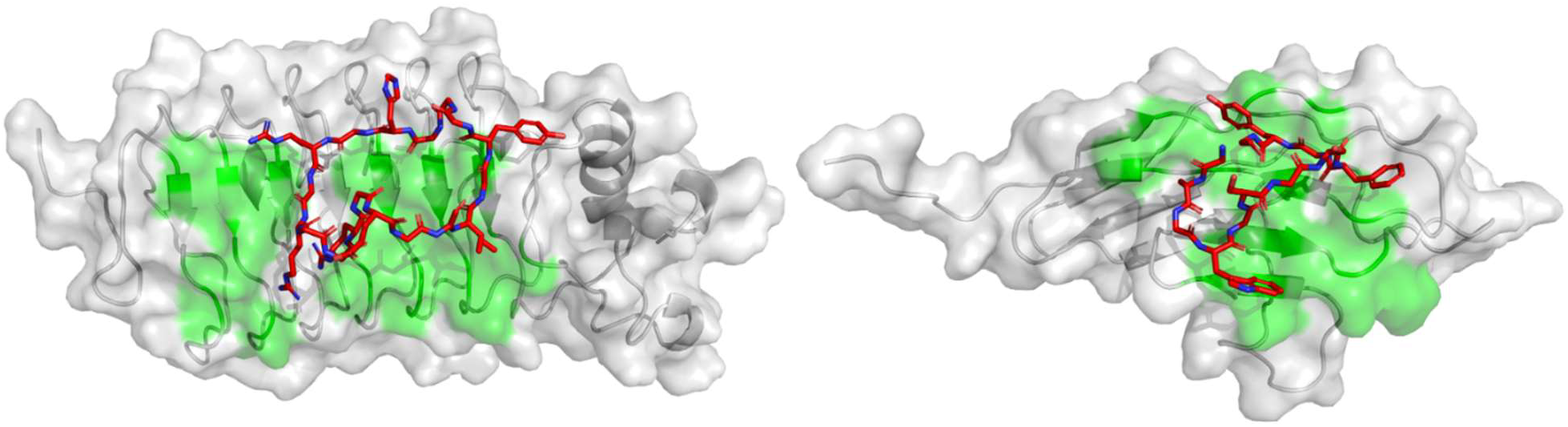

## 1. Introduction

Slit guidance ligands (SLITs) are secreted glycoproteins that regulate cellular positioning and migration during development through interactions with roundabout (ROBO) receptors.^1^ In mammals, SLIT1, SLIT2, and SLIT3 engage the second leucine-rich repeat domain (D2) of SLIT with the Ig1 domain of ROBO1 and ROBO2.^2,3^ The SLIT2/ROBO signaling axis plays multifaceted roles in organogenesis, tissue homeostasis, and tumor progression. Beyond its established functions in cell migration, it influences proliferation, apoptosis, adhesion, and angiogenesis in both normal and malignant contexts. SLIT2/ROBO signaling also contributes to hepatic fibrosis via PI3K/Akt activation and modulates cytoskeletal organization through adaptor proteins recruited to the ROBO cytoplasmic domain, thereby regulating motility and growth.^3-5^

In the nervous system, SLIT2/ROBO controls axonal repulsion,^6,7^ neuronal migration,^8^ and axon guidance.^9^ In the immune system, SLITs attract neutrophils while repelling lymphocytes and dendritic cells,^10-12^ and suppress macropinocytosis and cytotoxic polarization in macrophages.^13^ In endothelial cells, SLIT2–ROBO activation promotes angiogenesis in retinal and bone tissues by guiding tip-cell migration and polarization.^14-16^ Functionally, SLIT2 often exhibits proangiogenic and proinvasive activity in tumors,^17^ enhancing cancer cell migration,^18-20^ metastasis,^21^ and therapeutic resistance,^22^ particularly in colorectal, pancreatic, and osteosarcoma models. Conversely, tumor-suppressive roles have been reported in lung and breast cancers,^23-25^ underscoring the pathway’s context-dependent functions. In glioblastoma (GBM), evidence remains conflicting, with both inhibitory^26-28^ and tumor-promoting^29,30^ effects described.

Recent work has revealed SLIT2/ROBO signaling as an immune evasion mechanism in the GBM tumor microenvironment (TME).^31^ Elevated SLIT2 expression in both GBM patient samples and mouse models correlates with increased accumulation of immunosuppressive tumor-associated macrophages (TAMs) and vascular dysmorphia. Silencing SLIT2 in glioma cells or systemic inhibition using the SLIT2-trapping protein ROBO1-Fc prevented TAM polarization toward a tumor-supportive phenotype, reduced angiogenic gene expression, and improved tumor vascular function.^31^ These changes enhanced the therapeutic efficacy of both chemotherapy and immunotherapy in preclinical GBM models.^31^ Notably, SLIT2 blockade produced stronger effects on angiogenesis and T-cell response than prior TAM-targeting strategies within the GBM TME.^31^ Clinically, the only reported SLIT2/ROBO-targeted biotherapeutic, PF-06730512, was tested in patients with focal segmental glomerulosclerosis (NCT03448692) but was discontinued in 2023 due to insufficient efficacy at tolerated doses.^32,33^ Meanwhile, three active clinical trials (NCT03940820, NCT03941457, NCT03931720) are evaluating chimeric antigen receptor–natural killer (CAR-NK) cell therapies directed against ROBO1 in solid tumors.

Peptide therapeutics have emerged as an important class bridging the gap between small molecules and biologics.^34^ Among them, cyclic peptides stand out for their conformational rigidity, which enhances proteolytic stability, pharmacokinetic behavior, and binding affinity toward protein targets.^35^ In contrast to their linear counterparts, cyclic peptides can effectively engage shallow and extended interfaces typical of protein–protein interactions (PPIs), positioning them as promising scaffolds for targets traditionally considered “undruggable.”^36^ Over the past two decades, several cyclic peptides have gained clinical approval, validating this modality across oncology, infectious, and autoimmune diseases.^36^ Nevertheless, the rational design of cyclic peptides remains challenging due to their vast conformational landscape and the need for predictive methods that integrate both sequence and topological constraints. In this study, we present the first computational strategy for designing macrocyclic peptides that inhibit the SLIT2/ROBO1 interaction, followed by evaluation through TR-FRET–based inhibition assays and complementary biophysical validation. The most active candidates demonstrated favorable in vitro pharmacokinetic properties, including enhanced stability in simulated gastric and intestinal fluids and resistance to proteolytic degradation. Overall, this work paves the way for developing macrocyclic peptide therapeutics that selectively modulate the SLIT2/ROBO1 pathway, offering a foundation for future translational applications in cancer and neuroimmune disorders.

## 2. Materials and Methods

### 2.1. Computational design of macrocyclic peptides

The peptides were designed using the fragment-based approach Des3PI.^37^ Specifically, we used the advanced version Des3PI 2.0, a full description of which is presented in a forthcoming manuscript.^38^ Schematically, the 20 canonical amino acids are docked multiple times onto a targeted protein surface, then Des3PI 2.0 clusters the centers of mass of the side chains from all docking poses to identify high-density binding regions that we called hotspots. Finally, based on the amino acid composition within each hotspot, Des3PI generates possible macrocyclic peptide sequences, using glycine residues as linkers when hotspots are too far apart.

Des3PI 2.0 was applied independently to each of the two proteins of the SLIT2/ROBO1 complex (PDB ID: 2V9T).^39^ For each target protein, a docking box was defined to encompass all residues involved in the interactions with its partner. By default, Des3PI 2.0 outputs 20 macrocyclic peptide sequences for each application.

These 20 peptides were further ranked by using blind docking calculations (i.e. without predefining any binding area) onto their target protein, using AutoDock CrankPep (ADCP).^40^ Each docking pose yielded by ADCP was evaluated with a contact score representing the fraction of interfacial residues of SLIT2 or ROBO1 protein that interact with the docked peptide. A protein residue is considered in contact if it lies within 0.5 nm from the peptide. Finally, the ten peptides with the highest contact score among the top 5% of ADCP docking scores were proposed for synthesis and biological assays.

### 2.2. Synthesis of the peptides

See the Supplementary Material file.

### 2.3. SLIT2/ROBO1 TR-FRET assay

The assay was performed as we previously reported.^41^ See the Supplementary Material file for detailed experimental.

### 2.4. BLI validation of SLIT2/ROBO1 inhibition

See the Supplementary Material file.

### 2.5. Pharmacokinetic profiling

The assays for determining pharmacokinetic profile were performed as previously described.^42^

- Stability in intestinal simulated fluids (t1/2, min) was evaluated using erythromycin as a reference compound, t1/2, min = 453 min.^42^
- Stability in rat liver microsomes was evaluated using verapamil as a reference compound, t1/2, min = 6.3 min.^42^
- Human plasma stability was evaluated using propantheline as a reference compound, t1/2, min = 2.6 min.^42^

## 3. Results

### 3.1. Macrocyclic peptide sequences generated by Des3PI 2.0

The application of Des3PI 2.0 to SLIT2 identified seven hotspots and twenty peptide sequences (Figure 1A) occupying a large central region of the interaction area. Among these hotspots, three are predominantly composed of aromatic or hydrophobic amino acids and located close to several SLIT2 hydrophobic residues. For example, the red hotspot (Ile or Trp) is in contact with protein Leu378 and Tyr356, while the blue hotspot (Tyr or Trp) is at proximity to His406 and Tyr404. The yellow hotspot mainly contains the polar Ser amino acid which likely interacts with SLIT2 Ser332 or Tyr356 hydroxyl group. However, in three peptides out of 20, an Arg is found at this hotspot, which may be explained by the proximity of two negatively charged residues (Asp330 and Glu308) of SLIT2 protein. These residues also probably account for the strong presence of a Lys amino acid in the brown hotspot, located nearby. Finally, an Arg amino acid is dominantly encountered in the purple and orange hotspots, consistently with the presence of a highly polar pocket (Asn310, Asn333, Asn334, Thr311, and Gln309) at this area of the SLIT2 interface. Overall, the designed peptides could bind both the extensive hydrophobic region of the SLIT2 interface and the adjacent polar and charged areas.

**Figure 1.**
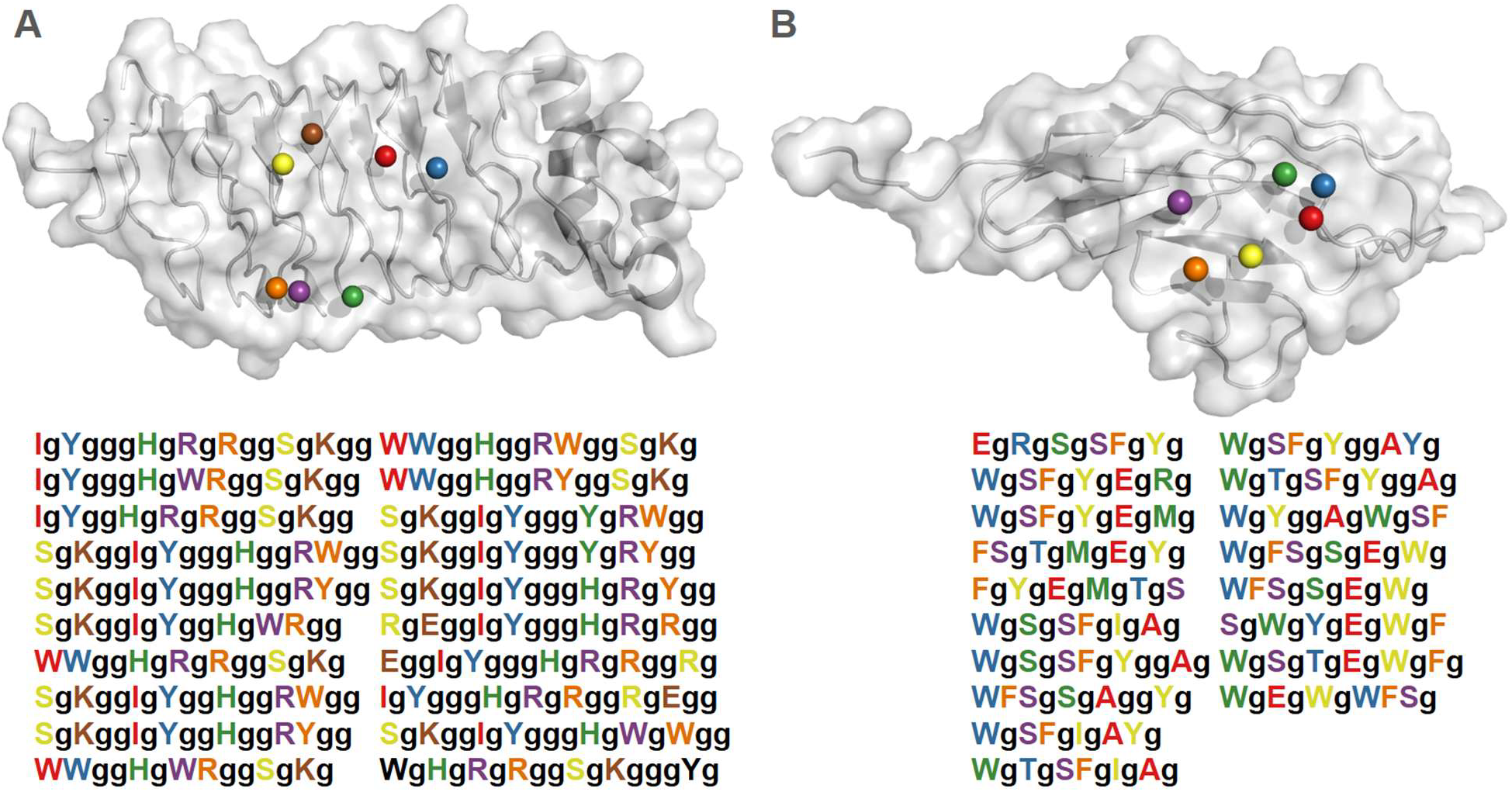
Hotspots and best sequences computed by Des3PI 2.0 on (A) SLIT2 and (B) ROBO1 target proteins. Bold lowercase ‘**g**’ represents glycine linker.

Regarding the ROBO1 interface, Des3PI 2.0 generated 18 peptide sequences having six hotspots (Figure 1B). As observed for SLIT2, a core of three aromatic hotspots can be identified. The yellow hotspot (composed almost exclusively of Trp or Tyr) is located near ROBO1 Leu127 and Phe128 residues, suggesting potential hydrophobic interactions. The blue and green hotspots are also mainly hydrophobic, being close to the protein Leu77, Val79, and Ala85 residues, although both can occasionally feature an Arg amino acid, possibly making a salt bridge with the close ROBO 1 Asp76 residue. Notably, the red hotspot exhibits a strong representation of the negatively charged Glu residue, which can be explained by the proximity of Arg131 on ROBO1 targeted surface. Similarly to SLIT2, the ROBO1-derived peptides contain hydrophobic amino acids essential for targeting the extensive hydrophobic region of the interface, complemented by polar or charged residues that may contribute to binding specificity.

The peptides generated by Des3PI 2.0 were submitted to blind docking calculations onto their respective protein target using ADCP, to rank them according to their ability to bind the targeted surface (quantified by their contact scores). Within the top 5% of ADCP scores, the 10 peptides with the highest contact scores are listed in Table 1. For the best peptide of each target protein, the binding mode retrieved by the blind docking calculations are shown in Figures 2A and 2B.

**Table 1:**
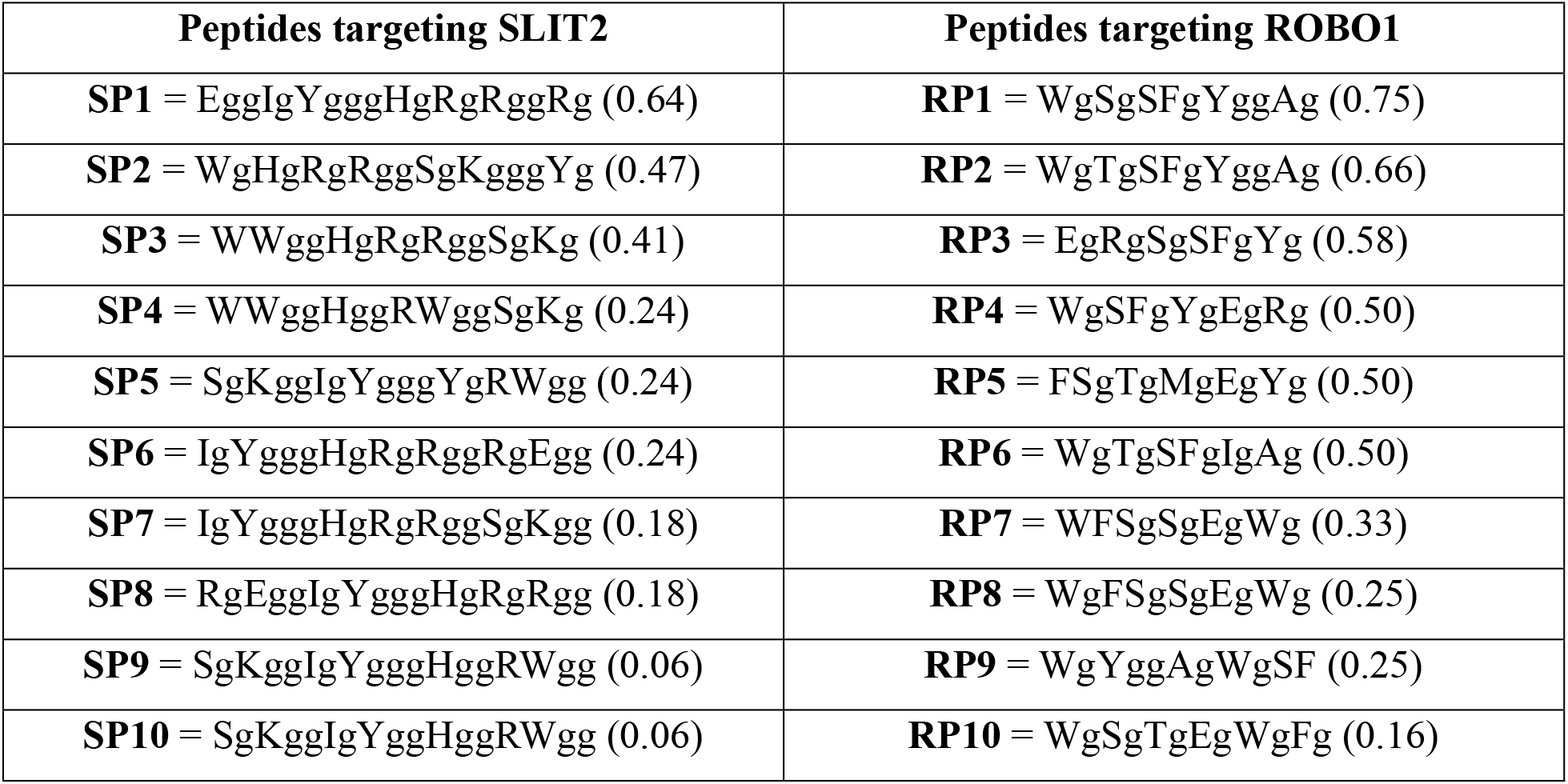
Name and sequence of the 10 best macrocyclic peptides designed by Des3PI 2.0 to target SLIT2 and ROBO1 interfaces. The contact score for each peptide is indicated in brackets.

**Figure 2:**
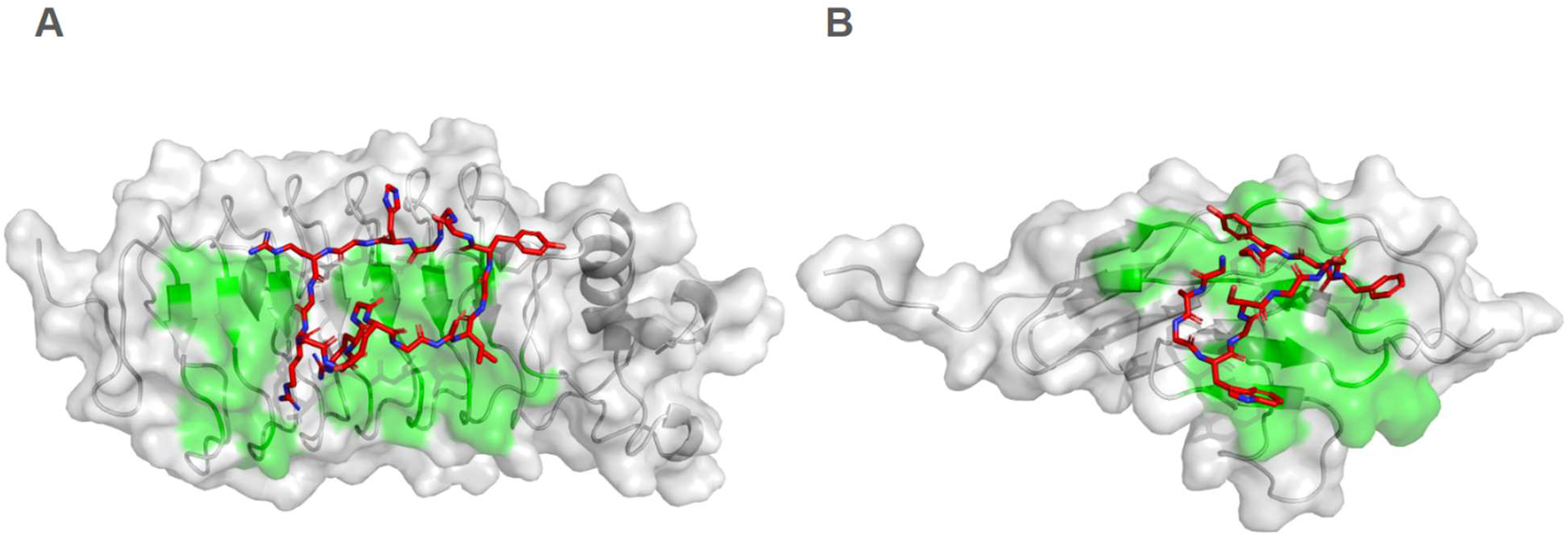
Binding mode of the best ranked macrocyclic peptide targeting **(A)** SLIT2 and **(B)** ROBO1. Residues of the protein interfaces are highlighted in green.

### 3.2. Primary screening by SLIT2/ROBO1 TR-FRET assay

Following computational design and ranking, the top four macrocyclic peptides targeting SLIT2 (**SP1**–**SP4**) and the top four targeting ROBO1 (**RP1**–**RP4**) were synthesized and experimentally evaluated for their ability to disrupt the SLIT2/ROBO1 interaction. The TR-FRET assay provided the first experimental evaluation of peptides designed to disrupt the SLIT2/ROBO1 interaction. This high-throughput and homogeneous format was chosen for its robustness and reproducibility in detecting weak-to-moderate affinity interactions typical of peptide–protein interfaces. Under the optimized conditions (5 nM SLIT2 and ROBO1, 1 h incubation), the assay exhibited high signal-to-background ratios and minimal vehicle variability, ensuring sensitivity to modest inhibitory effects at the single tested concentration of 200 µM.

As shown in Figure 3, the SLIT2-targeting peptides (**SP1**–**SP4**) produced substantially higher inhibition than the ROBO1-targeting series (**RP1**–**RP4**). Among the **SP** peptides, **SP4** demonstrated the strongest inhibition (62 ± 4%), followed by **SP3** (46 ± 3%), indicating effective disruption of the SLIT2/ROBO1 binding interface. **SP1** and **SP2** showed only weak inhibition (6–13%), suggesting suboptimal engagement or unfavorable conformations at the interface. In contrast, most ROBO1-targeting peptides exhibited minimal inhibition (<10%), except for **RP3**, which yielded a measurable reduction in TR-FRET signal (35 ± 3%), supporting a potential alternative binding orientation on the ROBO1 surface.

**Figure 3.**
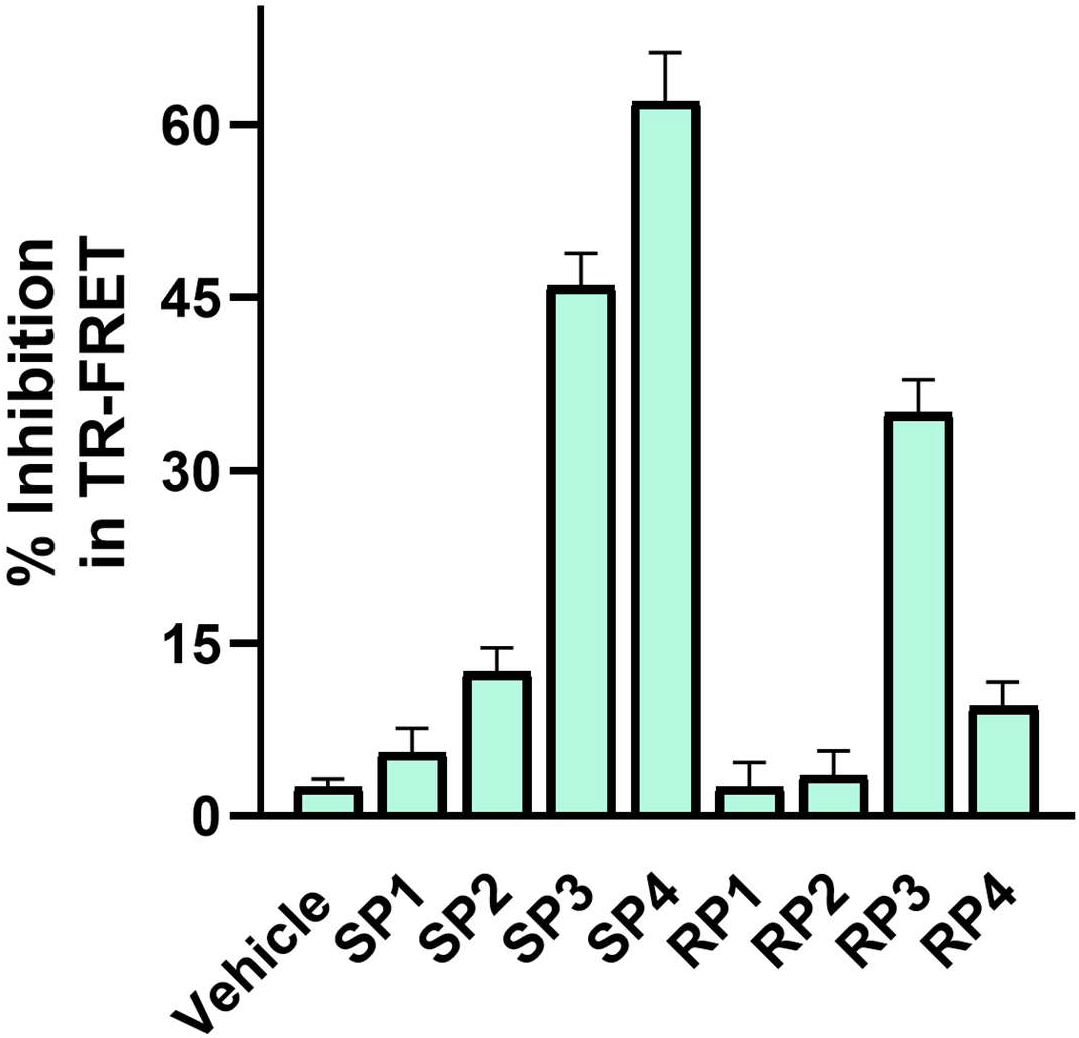
Inhibition of the SLIT2/ROBO1 interaction by macrocyclic peptides measured using the optimized TR-FRET assay. Each peptide (200 µM, n = 3) was evaluated for its ability to disrupt SLIT2/ROBO1 binding. Data are presented as mean ± SD. Among the SLIT2-targeting peptides, **SP4** and **SP3** exhibited the highest inhibition (≈ 60% and 45%, respectively), followed by weaker activity from **SP1** and **SP2**. The ROBO1-targeting peptides showed comparatively lower inhibition, with **RP3** producing ∼35% reduction in FRET signal.

Overall, the experimental inhibition supports the validity of the computational approach used for designing macrocyclic peptide inhibitors of SLIT2/ROBO1 interaction. Moreover, the lower activity observed for several peptides despite favorable in silico scores underscores the importance of experimental screening in capturing factors such as conformational flexibility, solubility, and aggregation, which cannot be fully predicted computationally.

These findings collectively confirm that macrocyclic peptides can directly interfere with the SLIT2/ROBO1 interaction, providing the first proof-of-concept for this therapeutic approach. The observed inhibition patterns also highlight SLIT2’s ligand surface as the more tractable site for peptide-based disruption, consistent with previous structural evidence showing that SLIT2 D2 contributes the majority of buried surface area in the SLIT2/ROBO1 complex. The TR-FRET results thus guided the selection of **SP3, SP4**, and **RP3** for further validation by biolayer interferometry (BLI) to quantify binding affinities and confirm direct competitive inhibition.

In summary, the TR-FRET data establish the functional relevance of the designed macrocyclic peptides, validate the computational framework, and identify lead candidates capable of perturbing the SLIT2/ROBO1 interface. These outcomes lay a strong foundation for subsequent optimization and mechanistic studies aimed at developing macrocyclic peptide scaffolds with improved potency, stability, and translational potential.

### 3.3. BLI validation of SLIT2/ROBO1 inhibition

To confirm direct binding and quantify inhibitory potency, the three most active peptides identified by TR-FRET (**SP4, SP3**, and **RP3**) were evaluated using a competitive BLI assay. Human ROBO1–Fc was immobilized on Protein A biosensors and exposed to recombinant SLIT2 pre-incubated with increasing concentrations of each macrocyclic peptide. Unlike the fluorescence-based TR-FRET assay, BLI provides a label-free and real-time measurement of complex formation, enabling direct observation of peptide-induced disruption of SLIT2/ROBO1 binding.

All three peptides produced dose-dependent decreases in SLIT2 binding response, confirming direct competition at the protein–protein interface (Figure 4). The concentration-dependent reductions in wavelength shift yielded well-defined sigmoidal inhibition curves from which IC_50_ values could be derived. The calculated IC_50_ values were 41 ± 5.2 µM for **SP4** (Figure 4A), 68 ± 7.5 µM for **SP3** (Figure 4B), and 115 ± 13 µM for **RP3** (Figure 4C), establishing the same potency trend observed in TR-FRET (**SP4** > **SP3** > **RP3**). The quantitative correlation between TR-FRET and BLI results validates the computational design and demonstrates that the observed inhibition arises from true disruption of SLIT2/ROBO1 complex formation rather than assay artifacts.

**Figure 4.**
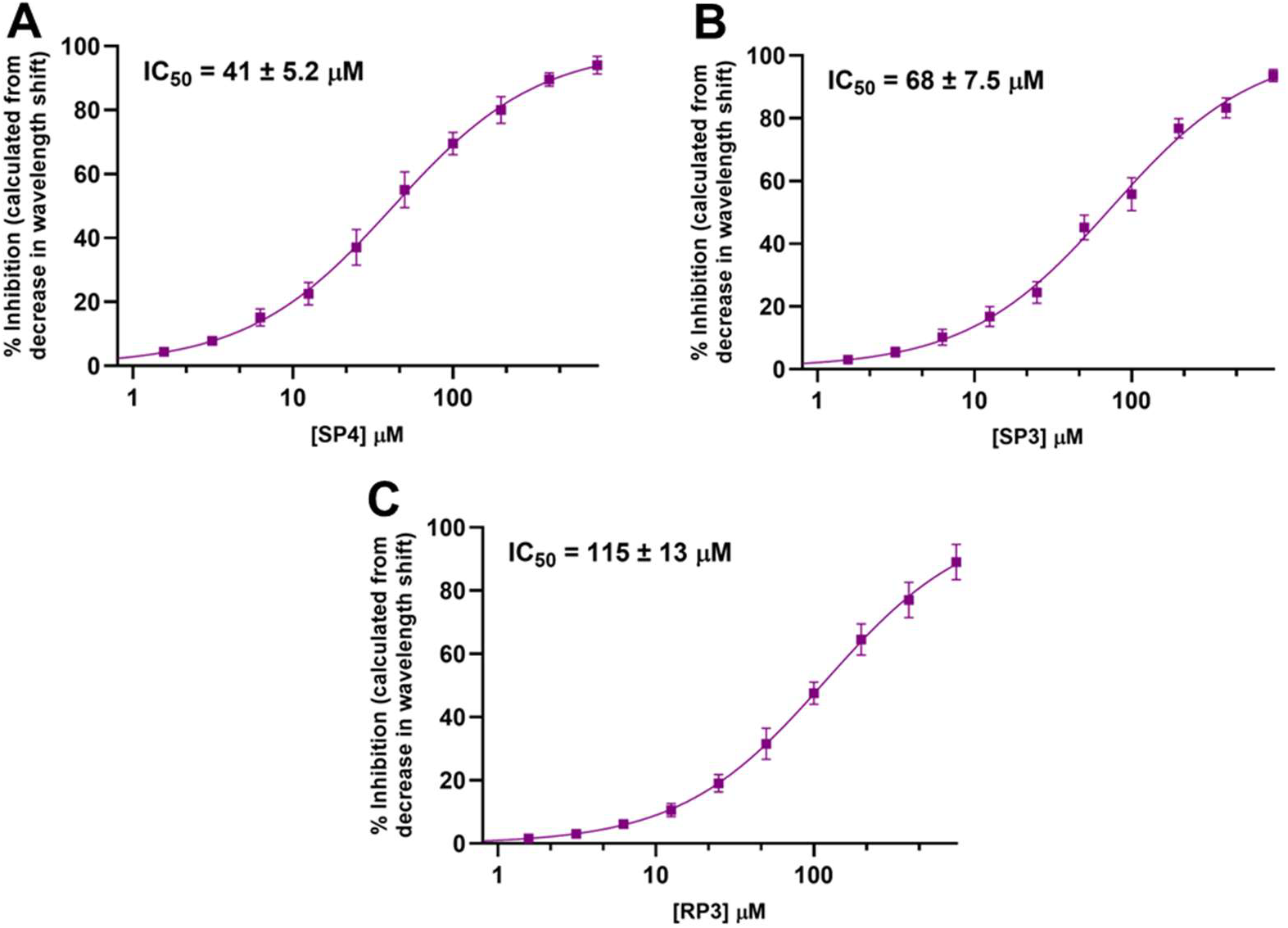
Dose-dependent inhibition of SLIT2/ROBO1 binding by macrocyclic peptides measured by biolayer interferometry (BLI). Human ROBO1–Fc was immobilized on Protein A biosensors and exposed to recombinant SLIT2 preincubated with increasing concentrations of each peptide. Shown are normalized inhibition curves for the top three peptides (**SP4, SP3**, and **RP3**) fitted with a four-parameter logistic model. Y-axis values represent % inhibition of SLIT2–ROBO1 binding, calculated from the relative decrease in BLI wavelength shift (Δλ) compared with the vehicle control (Δλ_0_). Data are shown as mean ± SD (n = 3).

Transitioning from TR-FRET to BLI was critical for confirming the mechanism of action. While TR-FRET efficiently identifies active compounds in a high-throughput format, BLI offers orthogonal verification under physiologically relevant, label-free conditions and provides quantitative IC_50_ values that guide structure–activity optimization. Together, these complementary assays establish the first evidence that computationally designed macrocyclic peptides can bind and inhibit the SLIT2/ROBO1 interface, highlighting **SP4** and **SP3** as promising scaffolds for further kinetic and structural characterization.

### 3.4. In vitro PK profiling

To gain early insight into the developability of the lead cyclic peptide **SP4**, we conducted an in vitro pharmacokinetic (PK) assessment encompassing chemical stability, lipophilicity, membrane permeability, and metabolic turnover. Such profiling is essential to evaluate the suitability of peptide scaffolds for further optimization, as their physicochemical and metabolic characteristics largely determine bioavailability and systemic exposure.

As summarized in Table 2, **SP4** exhibited moderate stability in simulated gastric fluid (t_1_/_2_ = 1.8 h) and remained notably more stable under simulated intestinal conditions (t_1_/_2_ = 7.6 h), consistent with enhanced backbone protection afforded by cyclization at neutral pH. In human plasma, **SP4** retained 89.4% of its integrity after 1 h, reflecting good resistance to proteolytic degradation. The measured LogD_7_._4_ value of 0.86 indicates a moderately hydrophilic profile favorable for systemic administration, while Caco-2 permeability assays revealed low-to-moderate bidirectional transport (Papp A→B = 1.78 × 10^-6^ cm/s; Papp B→A = 2.15 × 10^-6^ cm/s; efflux ratio ≈ 1.2), typical of macrocyclic peptides. In rat liver microsomes, **SP4** demonstrated moderate metabolic stability (t_1_/_2_ = 4.7 h), suggesting limited susceptibility to hepatic metabolism.

**Table 2.**
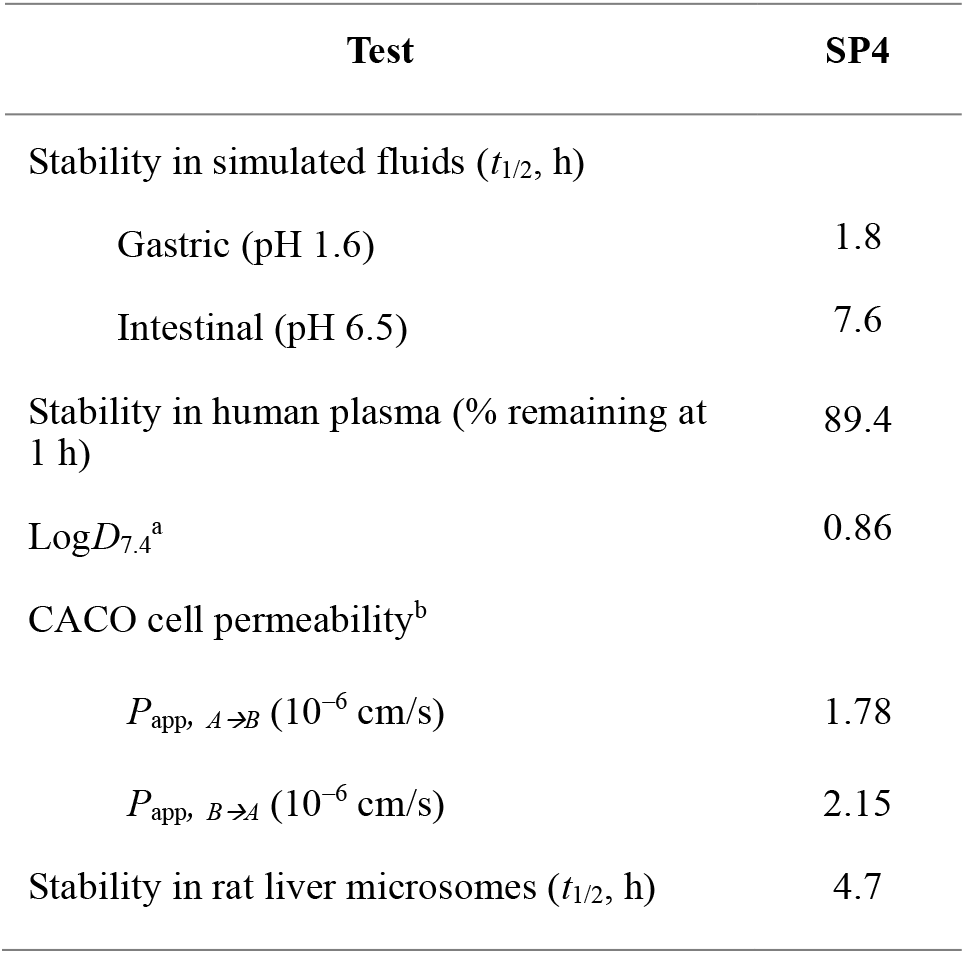
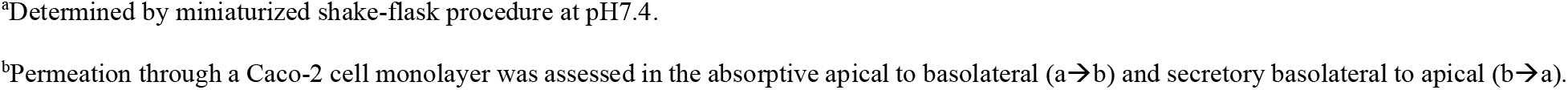
*In vitro* pharmacokinetic profile of **SP4**.

| Test | SP4 |
| --- | --- |
| Stability in simulated fluids ($t_{1/2}$, h) | |
| Gastric (pH 1.6) | 1.8 |
| Intestinal (pH 6.5) | 7.6 |
| Stability in human plasma (% remaining at 1 h) | 89.4 |
| $\text{LogD}_{7.4}^{\text{a}}$ | 0.86 |
| CACO cell permeability<sup>b</sup> |  |
| $P_{\text{app}} \text{ A} \rightarrow \text{B}$ ($10^{-6}$ cm/s) | 1.78 |
| $P_{\text{app}} \text{ B} \rightarrow \text{A}$ ($10^{-6}$ cm/s) | 2.15 |
| Stability in rat liver microsomes ($t_{1/2}$, h) | 4.7 |

<sup>a</sup>Determined by miniaturized shake-flask procedure at pH7.4.
<sup>b</sup>Permeation through a Caco-2 cell monolayer was assessed in the absorptive apical to basolateral (a→b) and secretory basolateral to apical (b→a).

Collectively, these findings indicate that **SP4** combines favorable plasma and intestinal stability with acceptable permeability and balanced lipophilicity, supporting its potential as a tractable macrocyclic scaffold for further chemical optimization and in vivo evaluation.

## 4. Conclusion

This study introduces a computational framework and biophysical validation strategy for the rational design of macrocyclic peptides targeting the SLIT2/ROBO1 interaction, an emerging axis implicated in tumor angiogenesis and immune suppression. By integrating interface-guided peptide generation with orthogonal TR-FRET and BLI screening, we identified **SP4** as a SLIT2– ROBO1 inhibitor exhibiting favorable in vitro stability and permeability profiles. The combined computational and experimental results validate the tractability of the SLIT2/ROBO1 interface and establish macrocyclic peptides as a viable modality for modulating this pathway. Looking forward, structure-guided optimization and in vivo evaluation of **SP4**-derived analogs could yield first-in-class therapeutics capable of reprogramming the SLIT2/ROBO1 axis in cancer and neuroimmune diseases, bridging a critical gap between target discovery and translational intervention.

## ACKNOWLEDGMENTS

We acknowledge funding support by R01CA293456 (PI Gabr) from the National Cancer Institute (NCI).

## Author Contributions

The manuscript was written through contributions of all authors. All authors have given approval to the final version of the manuscript.

