## Supporting Information for "Computational Design and Biophysical Validation of Macrocyclic Peptides as Inhibitors of SLIT2/ROBO1 interaction"

**Contents**

| **1.** | **Experimental** | **2** |
| --- | --- | --- |
| **2.** | **MS and HPLC data for the synthesized peptides** | **4** |

**1. Experimental:**

**1.1. Synthesis of the peptides:**

The peptides were synthesized on 2-CL resin, using standard Fmoc synthesis protocol with DIC/Cl-HOBt coupling, on an APEX 396 automatic synthesizer. The resin was swollen in DMF for 30 min, treated with 20v% Piperidine-DMF for 8 minutes at 50°C to remove the Fmoc protecting group and washed with DMF for three times. For the coupling reaction, the resin was added with Fmoc-protected amino acid, Cl-HOBt, DIC and NMP. The mixture was vortexed for 20 minutes at 50°C. Afterwards, the resin was washed with DMF once. The cycle of deprotection and coupling steps was repeated until the last amino acid residue was assembled. After the final Fmoc protecting group was removed, the resin was treated with 20v% acetic Anhydride-NMP for 20 minutes. The resin was then washed with DMF, DCM and dried with air. The peptides were cleaved using a TFA cocktail (95v%TFA, 2.5v%water and 2.5v%TIS) for three hours. Crude peptides were precipitated by adding ice-chilled anhydrous ethyl ether, washed with anhydrous ethyl ether three times, and freeze-dried. After head-to-tail cyclization, the crude peptides were loaded onto a prep-HPLC column and purified with a gradient of 10%-55%B within 45 minutes at a flow rate of 12 ml/min. The peptides were analyzed by LC-MS and confirmed to have >95% HPLC purity, and freeze-dried.

**1.2. SLIT2/ROBO1 TR-FRET assay**

Briefly, A time-resolved fluorescence resonance energy transfer (TR-FRET) assay was employed to evaluate the ability of cyclic peptides to disrupt the SLIT2/ROBO1 interaction. Recombinant human SLIT2 (His-tagged, Sino Biological) and the extracellular domain of ROBO1 (Fc-fusion, Sino Biological) were used as binding partners. The donor and acceptor fluorophores consisted of antihuman IgG Tb-conjugated antibody (donor) and anti-His d2-conjugated monoclonal antibody (acceptor) (Cisbio, PerkinElmer).

Assays were performed in 384-well white plates using PPI detection buffer (Cisbio) under optimized conditions of 5 nM SLIT2 and 5 nM ROBO1, with donor and acceptor concentrations of 0.25 nM and 2.5 nM, respectively. Plates were incubated for 1 h at room temperature before measurement on a Tecan Infinite M1000 Pro plate reader (excitation = 340 nm, emissions = 620 nm and 665 nm).

Each macrocyclic peptide was tested in triplicate at a final concentration of 200 µM in 0.1% DMSO. The TR-FRET signal was expressed as the ratio of emission at 665 nm to 620 nm × 100. Control wells included complete assay mixture with DMSO vehicle. Percent inhibition was calculated relative to the DMSO control, with decreased FRET signal indicating disruption of SLIT2–ROBO1 binding.

**1.3. BLI validation of SLIT2/ROBO1 inhibition:**

A competitive biolayer interferometry (BLI) assay was performed to quantify inhibition of the SLIT2/ROBO1 interaction by macrocyclic peptides. Human ROBO1 (hFc-tagged, Sino Biological) was immobilized on Protein A (ProA) biosensors and exposed to recombinant His-tagged human SLIT2 preincubated with increasing concentrations of each peptide. Measurements were acquired on an Octet RED96e instrument (Sartorius) in kinetics buffer (PBS, 0.02% Tween-20, 0.1% BSA, pH 7.4) at 25 °C with continuous shaking at 1,000 rpm. Each cycle included a baseline (60 s), association (120 s), and dissociation (180 s) phase. The response data were reference-subtracted and globally fitted using a 1:1 binding model to calculate inhibitory potency (IC₅₀). A dose-dependent reduction in SLIT2 binding signal confirmed that the macrocyclic peptides disrupted the SLIT2/ROBO1 interaction, providing orthogonal validation of the TR-FRET findings.

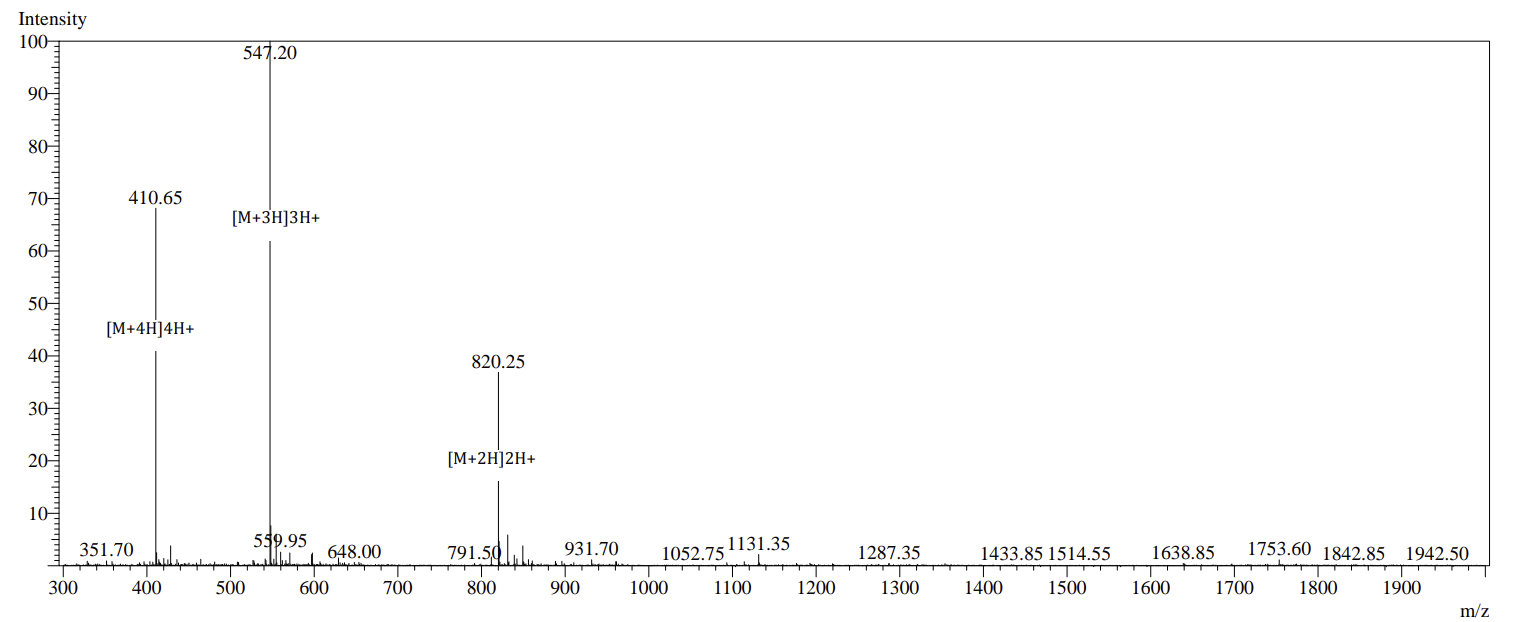
**Figure S1**. MS spectrum of **SP1**.

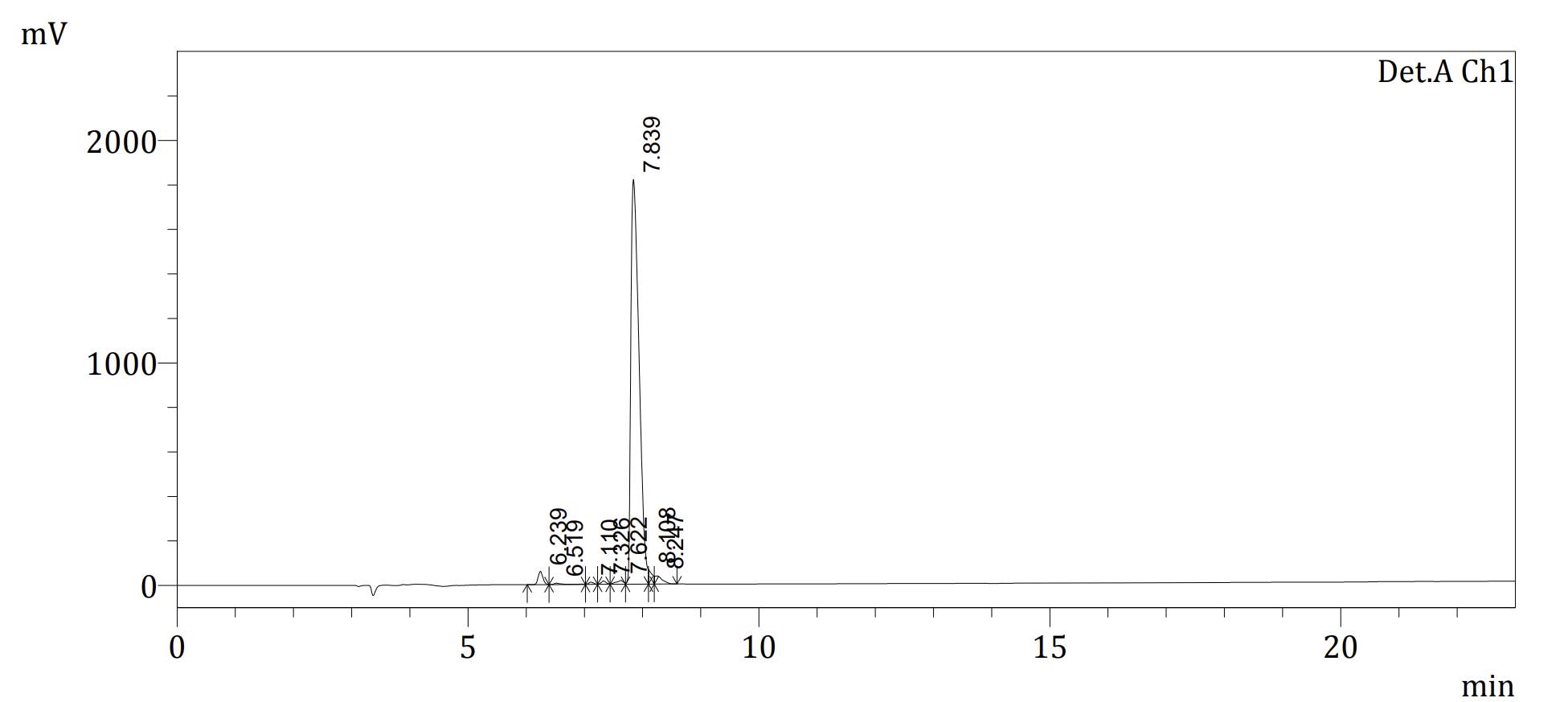

**Figure S2**. HPLC trace of **SP1**.

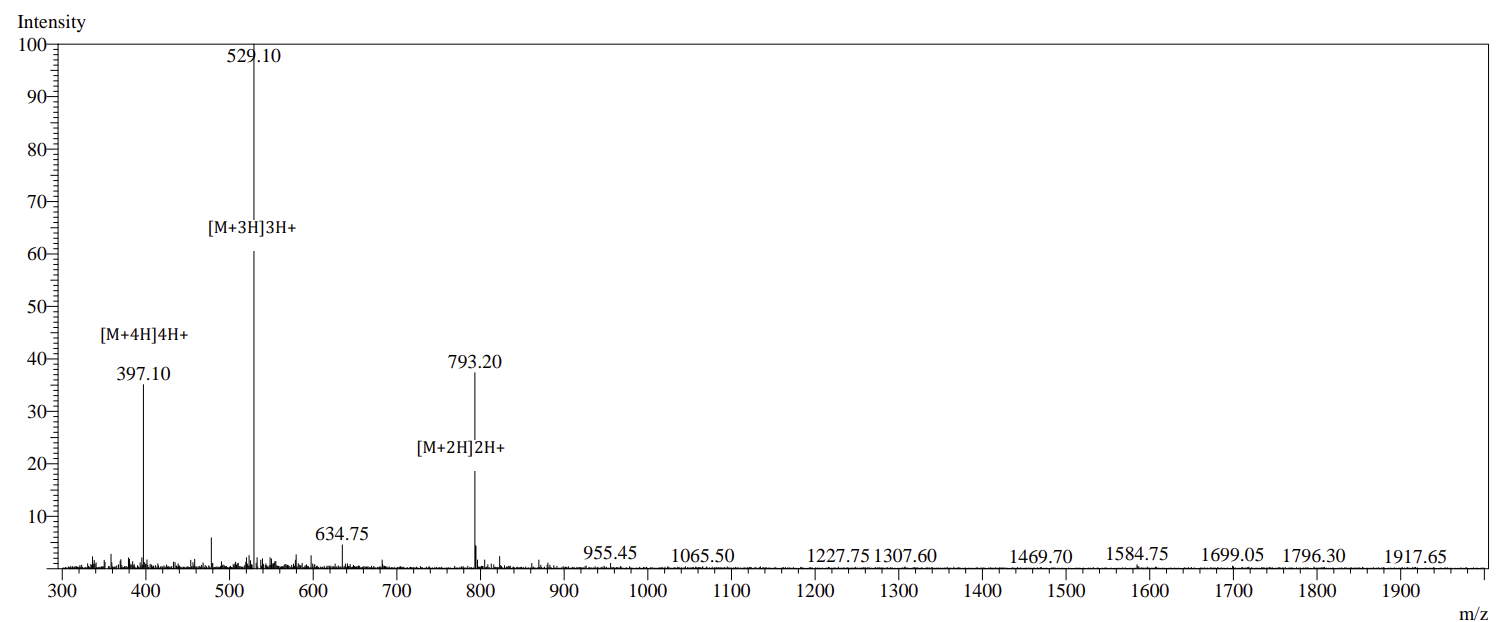

**Figure S3**. MS spectrum of **SP2**.

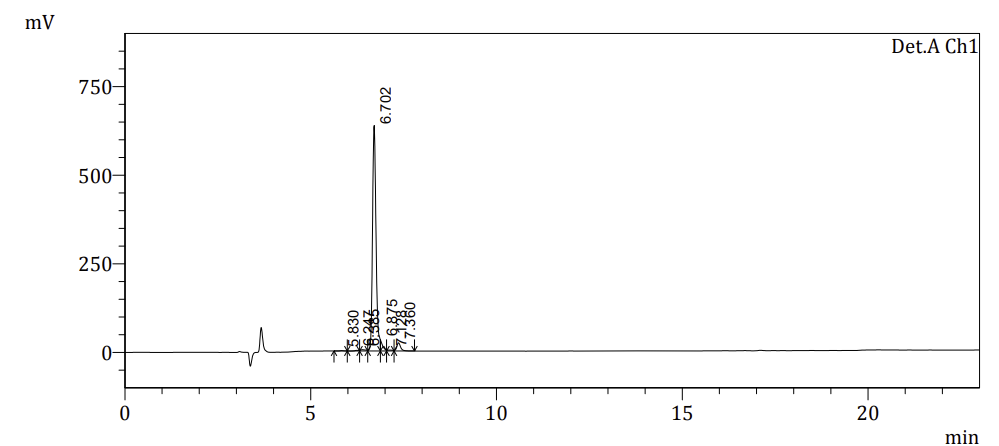

**Figure S4**. HPLC trace of **SP2**.

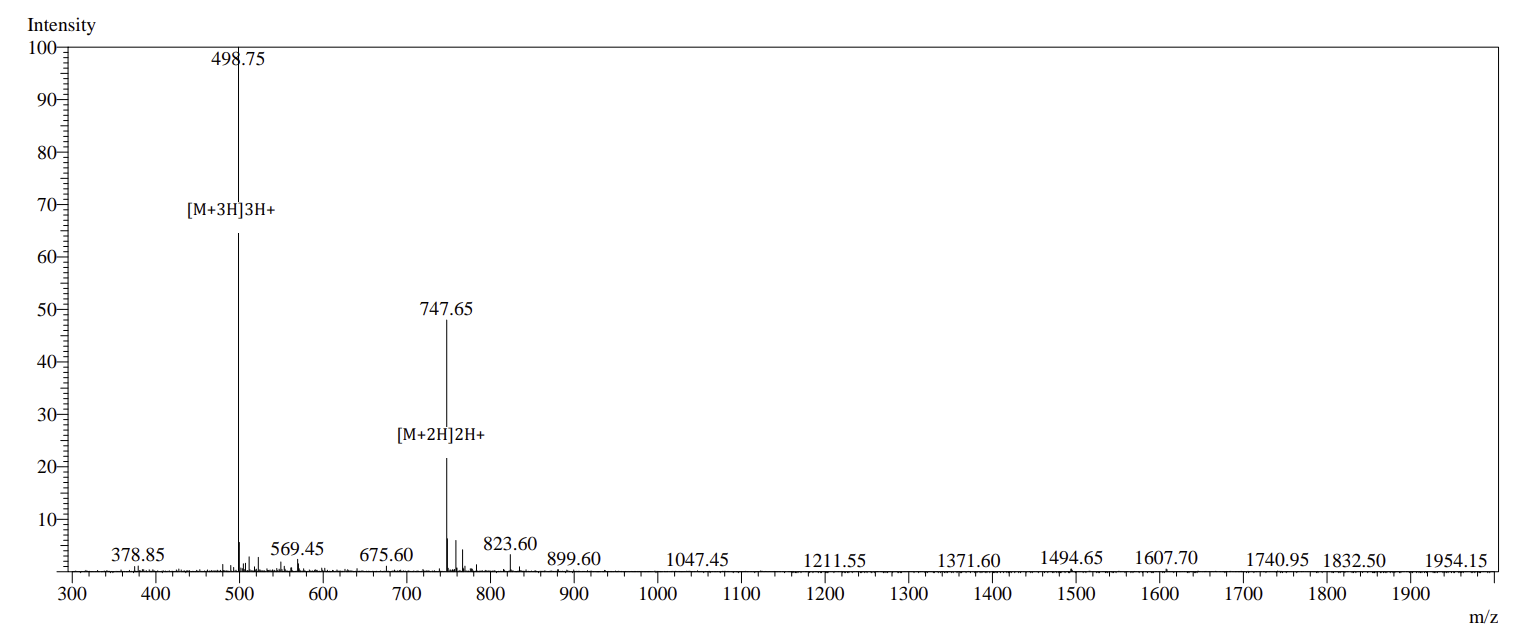

**Figure S5**. MS spectrum of **SP3**.

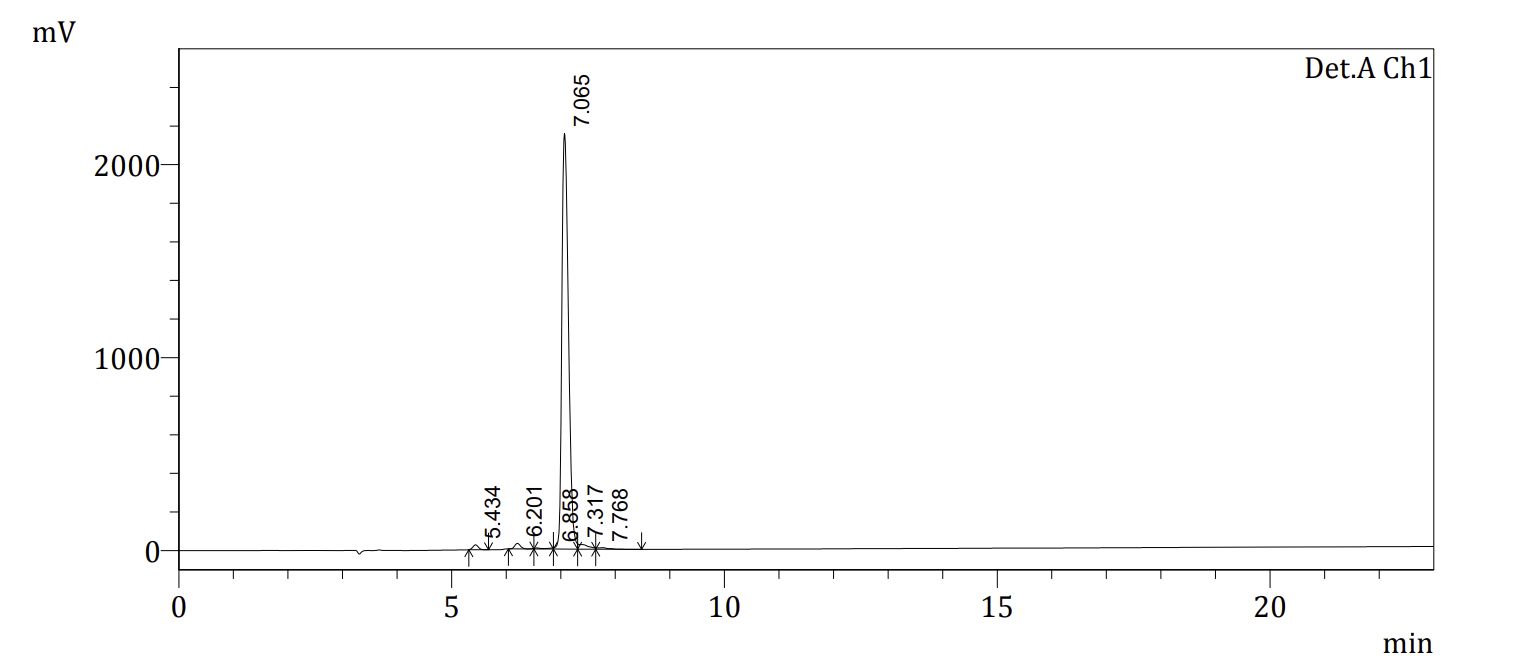

**Figure S6**. HPLC trace of **SP3**.

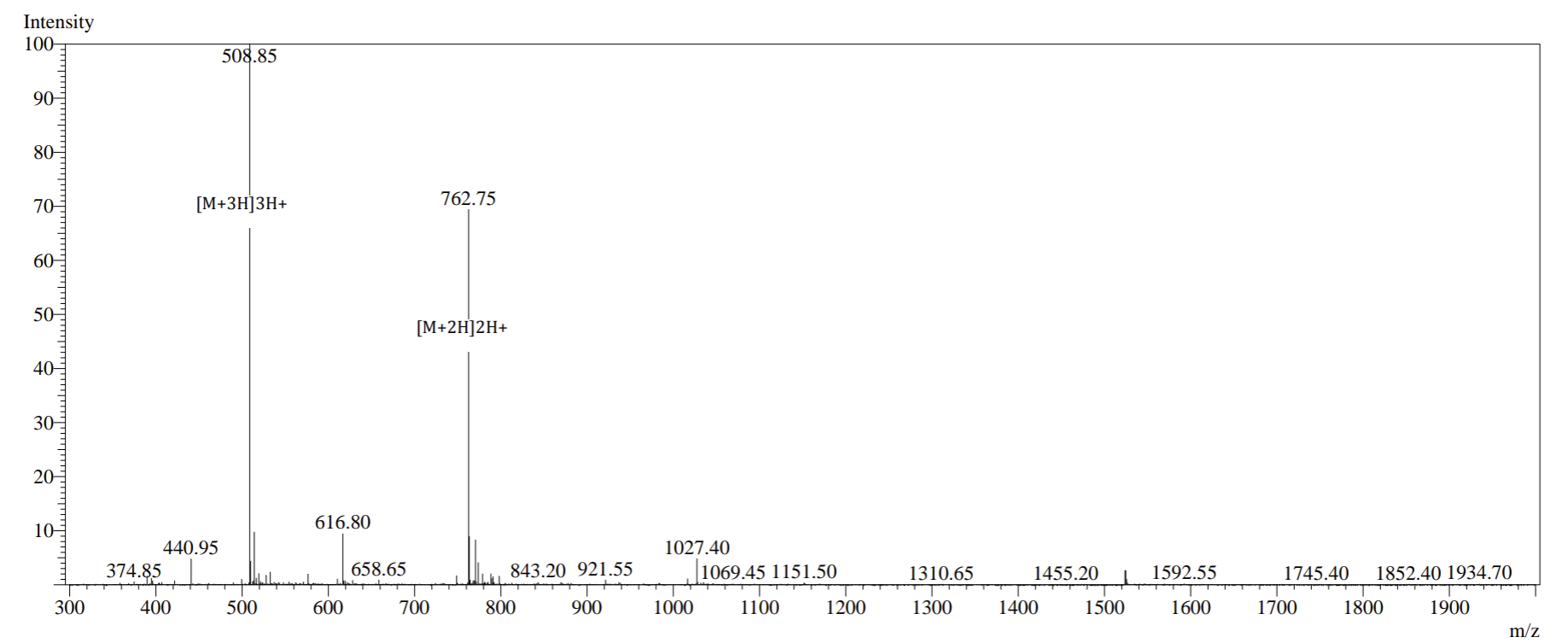

**Figure S7**. MS spectrum of **SP4**.

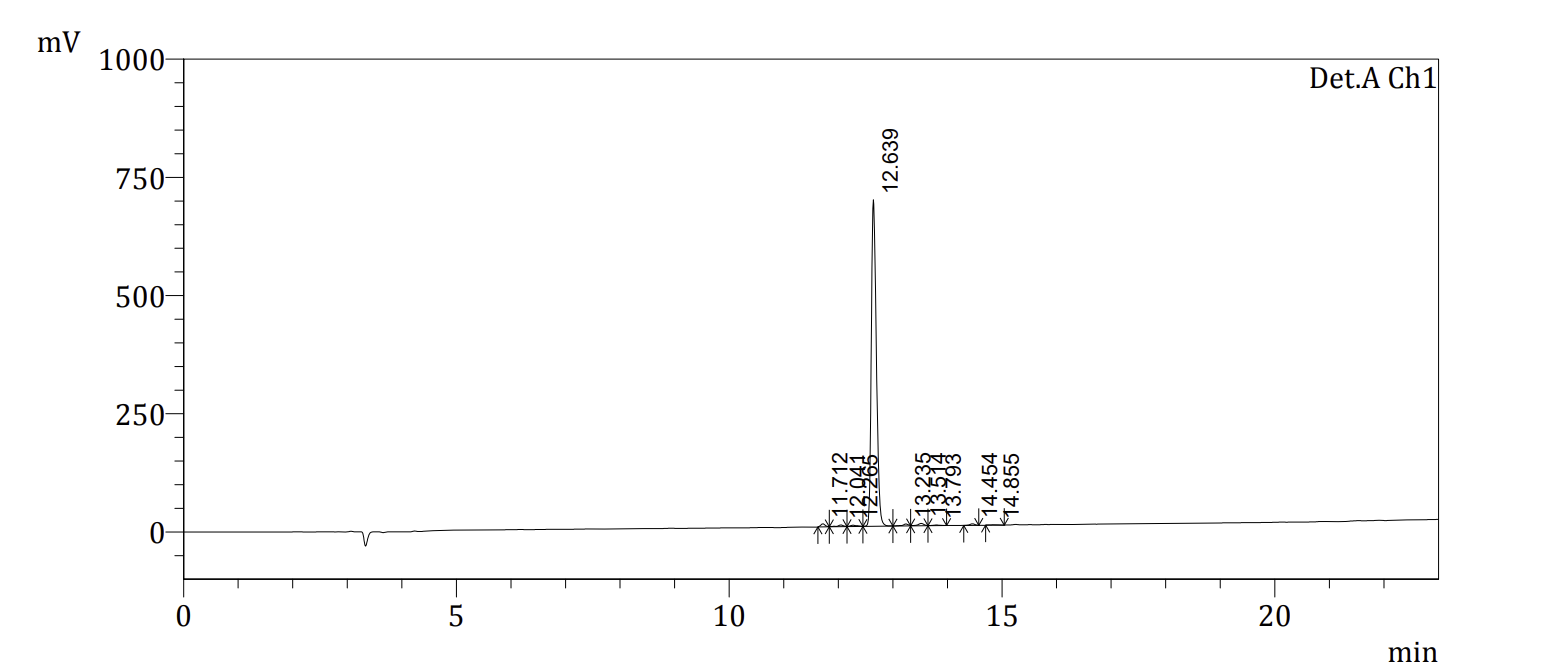

**Figure S8**. HPLC trace of **SP4**.

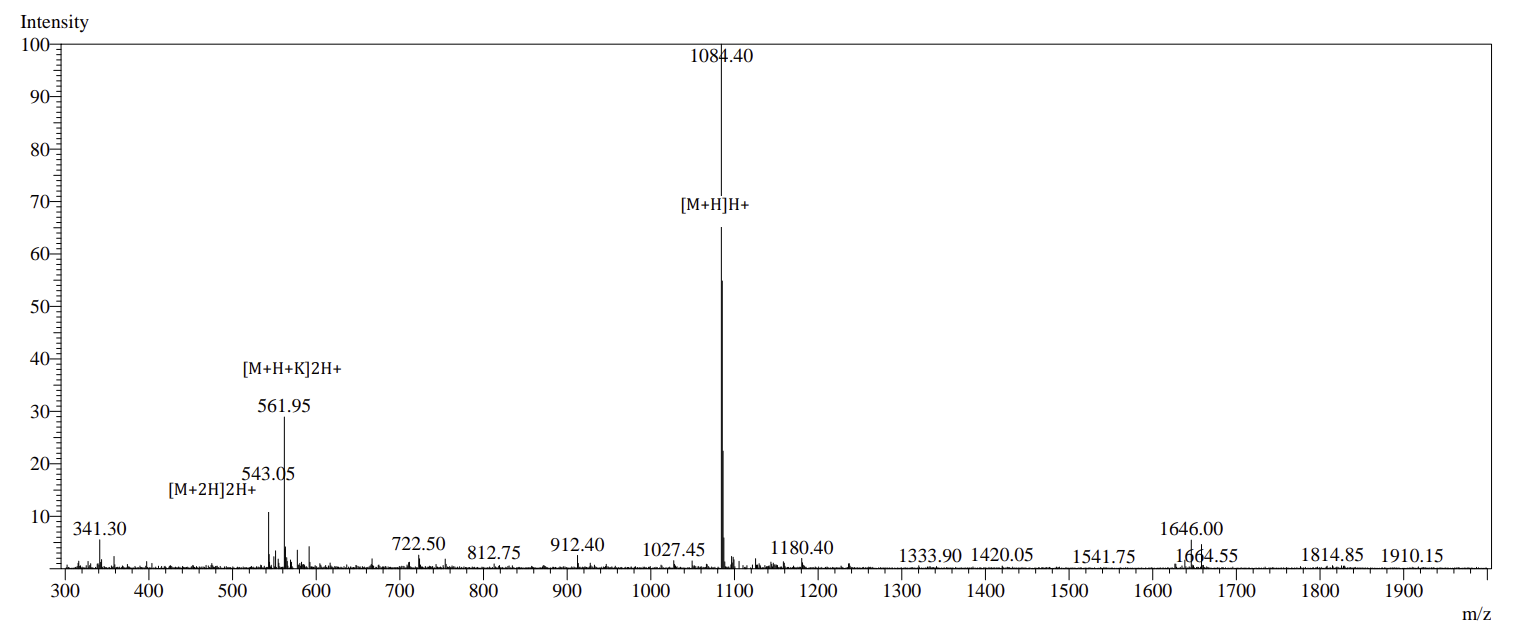

**Figure S9**. MS spectrum of **RP1**.

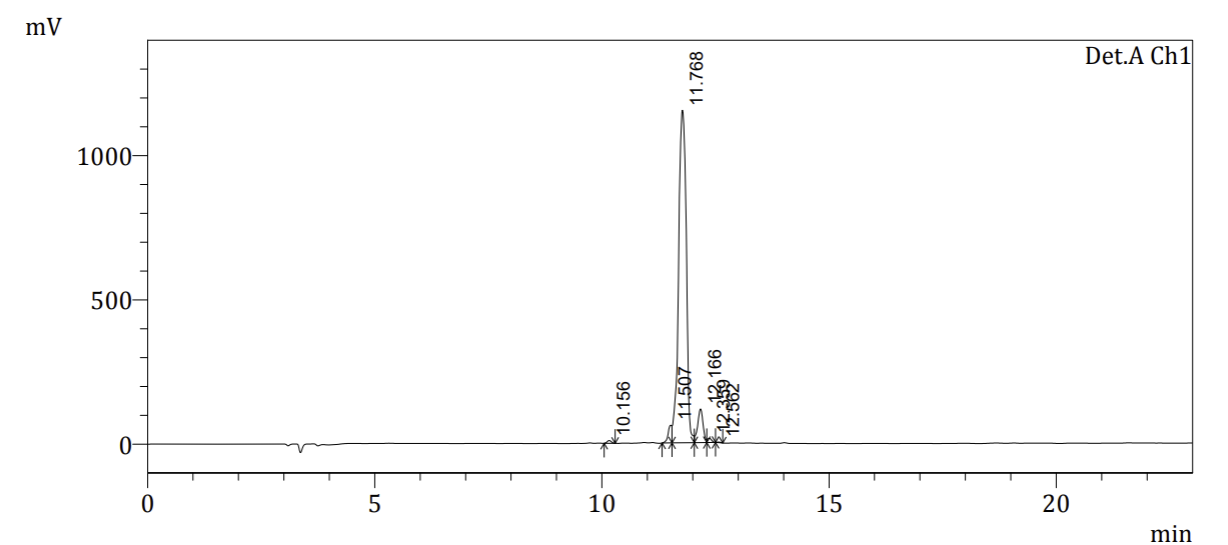

**Figure S10**. HPLC trace of **RP1**.

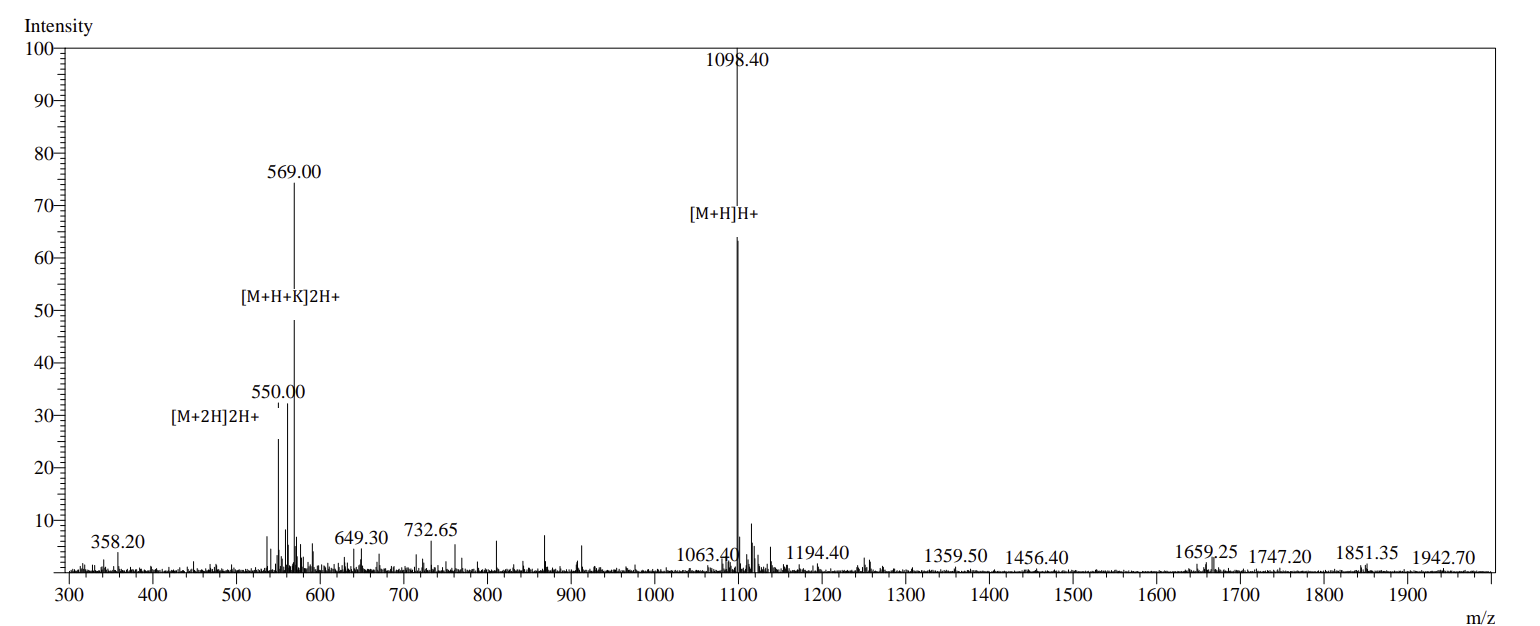

**Figure S11**. MS spectrum of **RP2**.

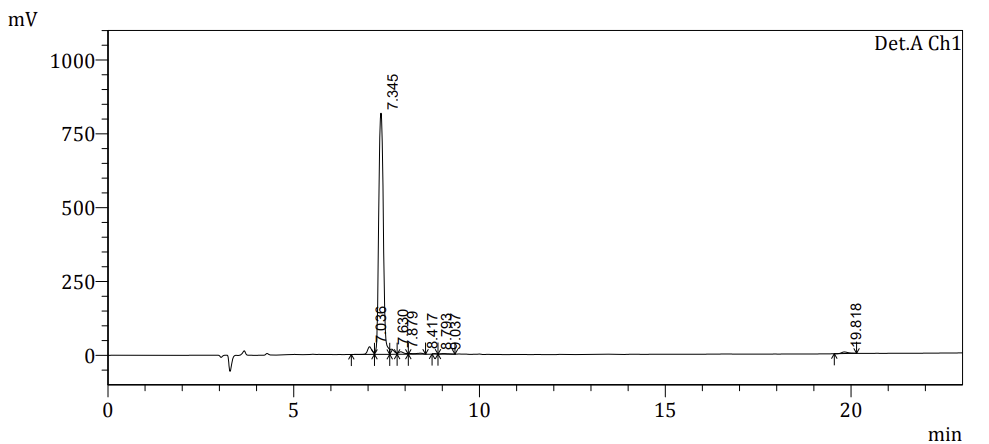

**Figure S12**. HPLC trace of **RP2**.

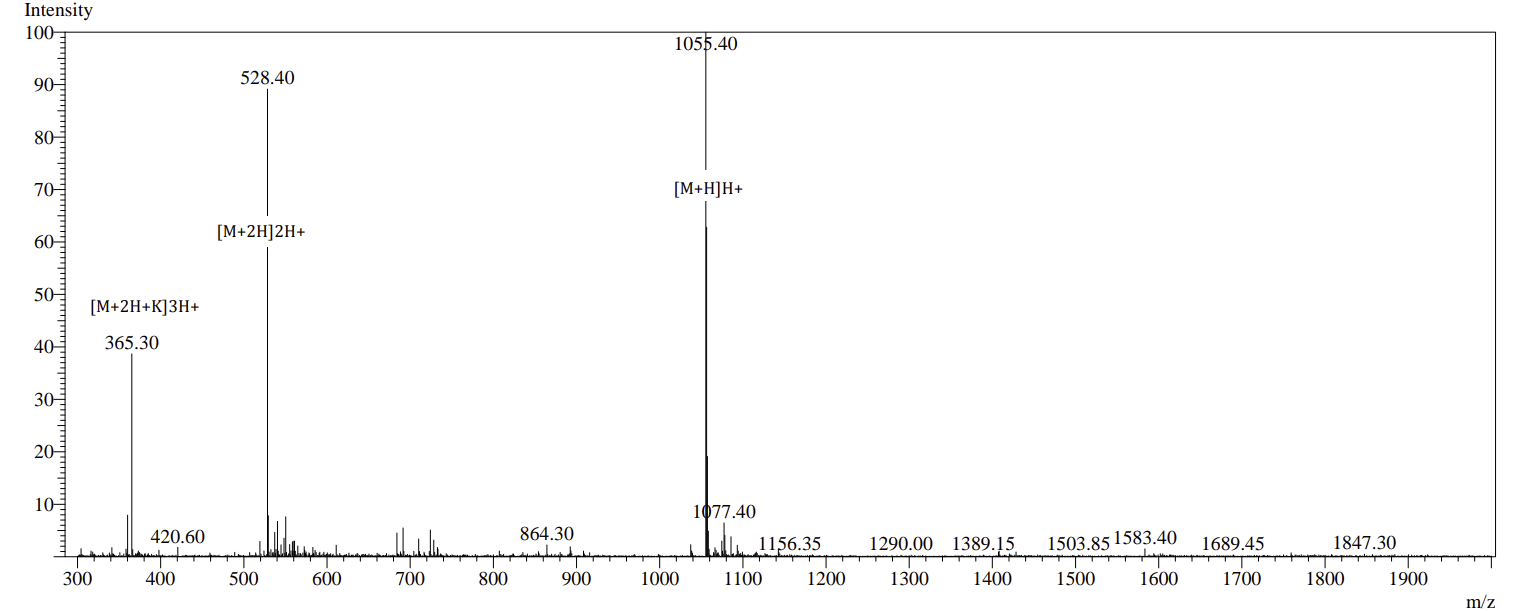

**Figure S13**. MS spectrum of **RP3**.

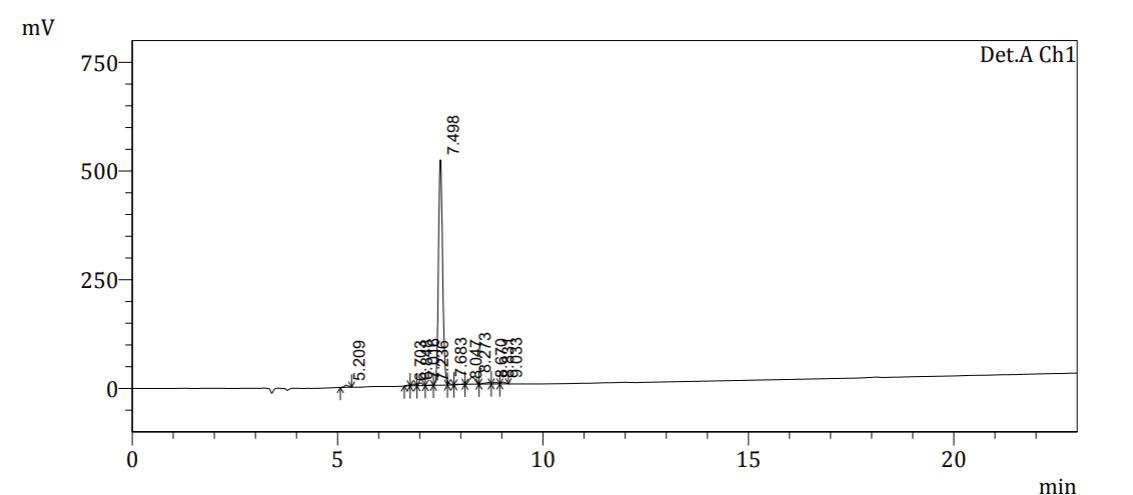

**Figure S14**. HPLC trace of **RP3**.

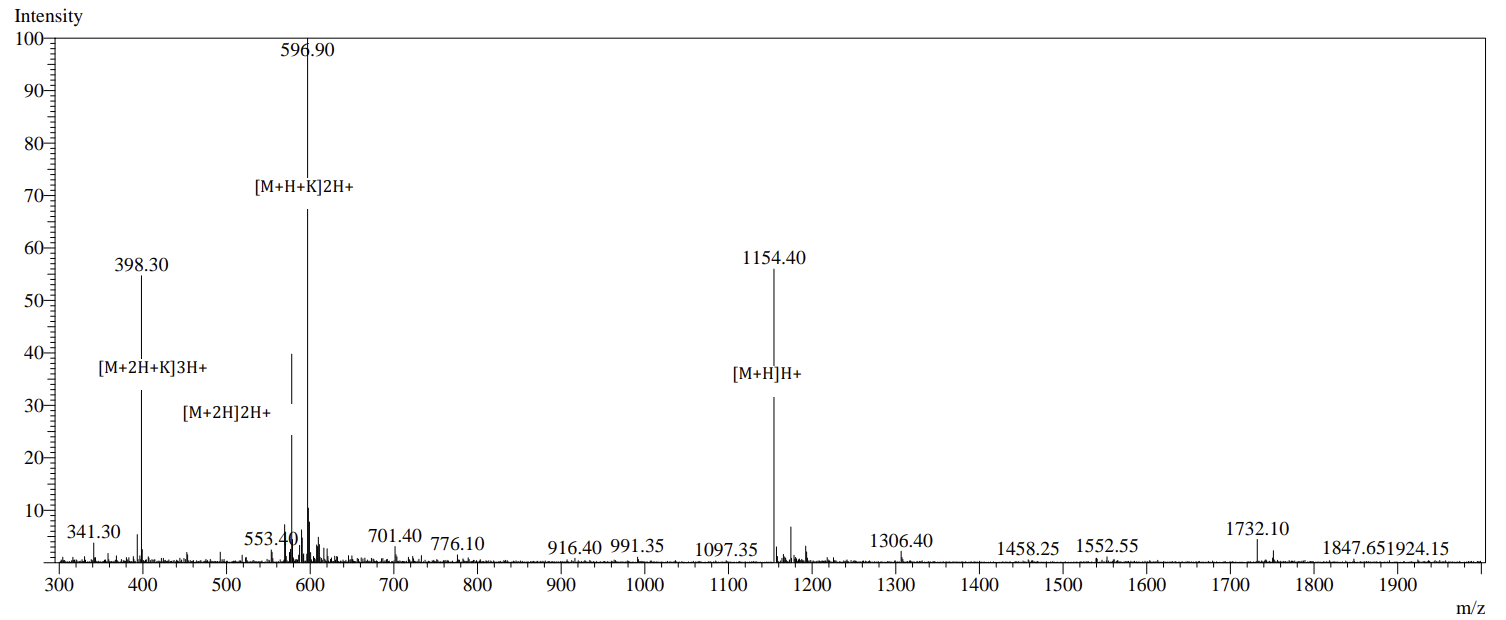

**Figure S15**. MS spectrum of **RP4**.

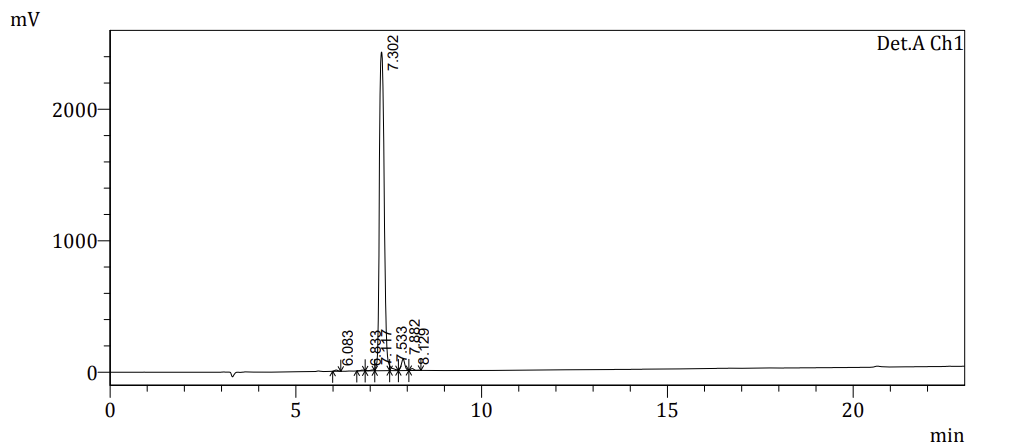

**Figure S16**. HPLC trace of **RP4**.
